# Ancestral sequence reconstruction of the Mic60 Mitofilin domain reveals residues supporting respiration in yeast

**DOI:** 10.1101/2024.04.26.591372

**Authors:** Friederike M. C. Benning, Tristan A. Bell, Tran H. Nguyen, Della Syau, Louise B. Connell, Yi-Ting Liao, Matthew P. Keating, Margaret Coughlin, Anja E. H. Nordstrom, Maria Ericsson, Corrie J. B. daCosta, Luke H. Chao

## Abstract

In eukaryotes, cellular respiration takes place in the cristae of mitochondria. The mitochondrial inner membrane protein Mic60, a core component of the mitochondrial contact site and cristae organizing system (MICOS), is crucial for the organization and stabilization of crista junctions and its associated functions. While the C-terminal Mitofilin domain of Mic60 is necessary for cellular respiration, the sequence determinants for this function have remained unclear. Here, we used ancestral sequence reconstruction to generate Mitofilin ancestors up to and including the last opisthokont common ancestor (LOCA). We found that yeast-lineage derived Mitofilin ancestors as far back as the LOCA rescue respiration. By comparing Mitofilin ancestors, we identified four residues sufficient to explain the respiratory difference between yeast- and animal-derived Mitofilin ancestors. Our results provide a foundation for investigating the conservation of Mic60-mediated cristae junction interactions.

## Introduction

Mitochondria are double membrane-defined eukaryotic organelles with essential metabolic and bioenergetic functions^1^. Cellular respiration, the process where adenosine triphosphate (ATP) is generated through oxidative phosphorylation, takes place in cristae, the characteristic invaginations of the inner mitochondrial membrane^2-6^ . Within the inner mitochondrial membrane, a neck-like membrane structure called the crista junction connects cristae to the inner boundary membrane and is an important site for transit of substrates essential for respiration and other physiological functions^7-9^.

Crista junction organization and stability is mediated by the mitochondrial contact site and crista organizing system (MICOS), a heterooligomeric inner mitochondrial membrane protein complex.^10-12^ MICOS consists of two subcomplexes that assemble independently and have distinct roles in crista junction formation and stability^13-15^. In yeast, the Mic60 subcomplex comprises Mic60 and Mic19 (and Mic25 in animals) and the Mic10 subcomplex consists of Mic10, Mic26, Mic27, and Mic12 (Mic13 in animals), which connects the two subcomplexes^14,16^.

Mic60 is the core shape-determining protein component of the Mic60 subcomplex. It is conserved across eukaryotes, Current addresses: Tristan A. Bell: Generate Biomedicines, Somerville, MA 02143, USA; Tran H. Nguyen: Medical Scientist Train-1 ing Program, Washington University in St. Louis, St. Louis, MO 63110, USA; Della Syau: Biological and Biomedical Sciences Program, Harvard Medical School, Boston, MA 02115 and all Mic60 proteins share a similar domain architecture with an N-terminal transmembrane helix followed by a large C-terminal region facing the intermembrane space^17^. Mic60 localizes at crista junctions, and together with Mic19, tethers the crista junction inner membrane region to the outer mitochondrial membrane via protein-protein interactions, including those with the sorting and assembly machinery for b barrel proteins (i.e., SAMs).^12,18-21^ Moreover, Mic60 is also an ancient MICOS component, with homologues identified in alphaproteobacteria, the extant cousins of a hypothesized endosymbiosed mitochondrial precursor^22^. Alphaproteobacterial Mic60 is found in intracytoplasmic membranes, which are functionally and morphologically analogous to mitochondrial cristae^17,23^. Bacterial and eukaryotic Mic60 share several functional and biophysical similarities, such as the ability to deform membranes^17,24^.

Downregulation of Mic60 causes the loss of crista junctions^11,12,18,25^, respiratory growth defects^26-29^, and is a hallmark of several neurodegenerative disorders such as Parkinson’s disease, Alzheimer’s disease, and various myopathies^30-32^. It has been established that Mic60’s most conserved region, its C-terminal Mitofilin domain, is important for respiration^27-29^ and interacts with Mic19, Mic10, and Mic13^33-35^. Recent structural work with truncation constructs of the Mic60 Mitofilin domain provide initial glimpses of its fold and interactions. A crystal structure of the region encompassing the lipid binding site 1 (LBS1) (a4 in Figure 1) fused to the three C-terminal helices (*α*6-8) of *Thermochaetoides thermophila* Mic60 has been determined. This construct was covalently linked to the C-terminal CHCH domain of *Thermo-chaetoides thermophila* Mic19^34^.

**Figure 1.**
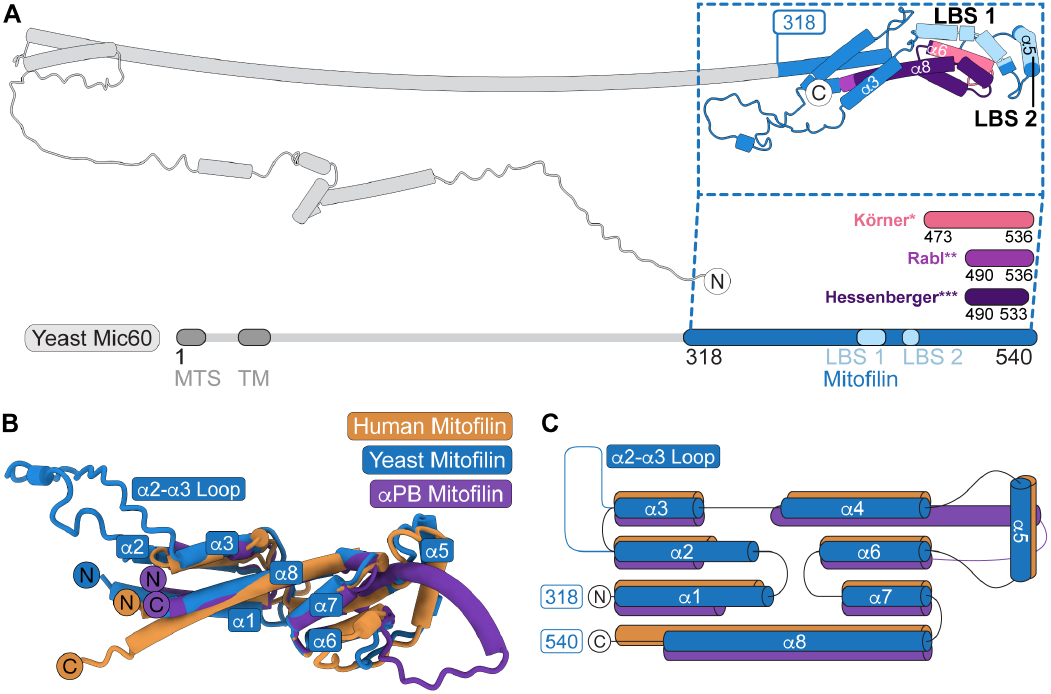
Boundaries of Mic60’s C-terminal Mitofilin domain defined based on its structural conservation. **(A)** AlphaFold2-predicted *S. cerevisiae* Mic60 structure (AF-P36112-F1) and linear Mic60 scheme showing different definitions of mitofilin domain boundaries. Blue (residues 318-540) denotes more generous domain boundaries based on structural features and alignments, which we used in all experiments in this study and which includes all other domain definitions. Previously published experimentally determined Mitofilin domain boundaries are colored in salmon (*, residues 473-536 by Körner *et al*.^42^), light purple (**, residues 490-536 by Rabl *et al*.^11^), and dark purple(***, residues 490-533 by Hessenberger *et al*.^35^). MTS: mitochondrial target sequence; TM: transmembrane region; LBS 1 and LBS 2: lipid binding sites 1 and 2 (light blue)^35^. **(B)** Superposition of AlphaFold2-predicted structures of the yeast (blue; residues 318-540), human (gold; residues 562-758) and al-phaproteobacterial (purple; residues 270-433) Mitofilin domains. RMSD by residue values for superposition on the yeast Mitofilin domain: 1.04 Å for human Mitofilin (Uniprot ID: Q16891) relative to yeast, 0.92 Å for al-phaproteobacterial Mitofilin (Uniprot ID: B9KRE6) relative to yeast. Sequences used for AlphaFold2 structure prediction are listed in Table S1. αPB: alphaproteobacterial. **(C)** Topology diagram of the human (gold), yeast (blue) and alphaproteobacterial (purple) Mitofilin domain. Domain boundaries for yeast Mic60 Mitofilin are listed next to the N and C terminus.

It remains unclear how Mic60’s Mitofilin sequence encodes its crista junction supported functions, in particular cellular respiration. While Mic60’s Mitofilin domain lipid binding sites have been investigated^35^, how self-assembly of full-length Mic60 and MICOS mediates cristae junction stability remains a major outstanding question. Furthermore, Mic60 coordinates an interaction network with respiratory complexes^36^ and other cristae-stabilizing proteins such as the SAM complex, Opa1 in animals, and the TOM complex^19,37-42^. In this respect, we lack detailed information on many aspects of Mic60 structure-function at the domain, protein, complex, and interaction-network levels.

As a step towards exploring Mic60’s sequence determinants supporting respiration, we leveraged the Mitofilin domain’s remarkable conservation across eukaryotes and performed ancestral sequence reconstruction, a molecular phylogenetic approach complementary to classical experimental structure determination^43^. First, using respiratory growth assays, we established that replacing the yeast Mitofilin domain with that of another yeast rescues cellular respiration. Next, we observed that substitution of the yeast Mitofilin domain with human or alphaproteobacterial homologues at the endogenous locus did not rescue respiration^24^. This finding, using CRISPR-edited lines, contrasts with previous plas-mid-based complementation. To explore how Mitofilin-supported respiration diverged between yeast and human, we used ancestral sequence reconstruction to reconstruct ancestral Mitofilin sequences and resurrected them for *in vivo* characterization. We found that common Mitofilin ancestors as far back as the putative last opisthokont common ancestor (LOCA) rescues mitochondrial respiration in yeast. Comparing reconstructed ancestors with different rescue phenotypes, we identified four residues in the Mitofilin fold that are involved in mitochondrial respiration. We also undertook mitochondrial ultrastructural and network morphology analyses, and describe a phenotype specific to a mutant of these four ASR-identified residues. Transmission electron microscopy (TEM) revealed that respiratory function may not strictly correlate with the presence of wildtype-like crista junctions. These findings, obtained from ancestral sequence reconstructions spanning ∼1 billion years of evolution^44^, pave the way for further studies investigating the timing of functional interactions in an ancient protein likely present during the initial eukaryogenesis event.

## Results

### A redefined domain boundary for the Mic60 Mitofilin domain

To address how Mic60’s C-terminal Mitofilin domain’s sequence encodes its respiratory function, we first revisited the definition of its domain boundaries. Despite extensive study, the definitions of Mic60’s Mitofilin domain boundary vary (colored in pink, light purple, and dark purple in Figure 1A). While it has been established that the Mitofilin domain extends to the C terminus, multiple N-terminal boundaries have been proposed (residues 473 and 490, respectively)^11,27,35,42,45^. We applied the protein-fold prediction tool AlphaFold to consider conserved structural elements to better define the Mitofilin N-terminal boundary^46^. Superimposing AlphaFold2 predictions of Mic60 from *H. sapiens, S. cerevisiae* and the alphaproteobacterium *C. sphaeroides*, we observed high structural conservation for a C-terminal region ranging from residues 318-540 (*S. cerevisiae* numbering; colored in blue in Figure 1A-C, Figure S1 for prediction confidence). This region includes all previously assigned domain boundaries for the Mitofilin domain but extends an additional three α helices in the N-terminal direction. These three additional helices are confidently predicted to form a 4-helix bundle with the C-terminal part of the previously defined Mitofilin domain (Figure 1A, Figure S1). It also includes the two previously proposed lipid binding sites (LBS 1 and LBS 2; residues 427-464; colored in light blue in Figure 1A)^11,35,42^ (Figure 1A). Our redefined Mitofilin domain is predicted to comprise two 4-helical bundles and an additional α helix, α5, which contains most residues of the second previously assigned lipid binding site (LBS 2; residues 454-464) and protrudes from the outer 4-helix bundle (Figure 1B, C). Residues of the other lipid binding site (LBS 1; residues 427-444) are within the fourth a helix (α4) of the outer 4-helix bundle. Helix α8 spans both helical bundles. Based on these structural alignments, we extended the Mitofilin domain boundaries to better reflect its conserved domain structure and used this larger Mitofilin region (residues 318-540) in all subsequent experiments.

### Alphaproteobacterial and human Mitofilin do not rescue cellular respiration in yeast

To test for Mic60 respiratory function, we applied an established respiratory growth assay in *S. cerevisiae* (Figure 2A). In this assay, oxidative phosphorylation-dependent respiratory growth was monitored using glycerol and ethanol (YPEG media; see Methods) as a non-fermentable carbon source. Only *S. cerevisiae* strains capable of mitochondria-dependent cellular respiration grow on YPEG. To control for mitochondria-independent growth deficits, we grew parallel cultures of all strains in dextrose (YPD media), a fermentable carbon source (Figure S2). We used CRISPR-Cas9 to generate Mic60 mutations at the endogenous locus in the genome of the S288C strain of *S. cerevisiae*. Consistent with previous work using derivatives of BY4741 *S. cerevisiae*^27,29,42^, we observed that knocking out Mic60 in *S. cerevisiae* (*mic60*Δ) leads to poor growth compared to wildtype under non-fermentable conditions (Figure 2A). Similarly, poor respiratory growth has been previously reported for C-terminally truncated versions of Mic60 (Mic60_1-140_, Mic60_1-472_, and Mic60_1-491_ all show a reduced growth phenotype in *S. cerevisiae*)^27,29,42^. As expected, our truncation based on our redefined Mitofilin domain definition (Mic60_1-317_; hereafter referred to as *mitofilin*Δ), also shows a reduced growth phenotype similar to *mic60*Δ (Figure 2A).

**Figure 2.**
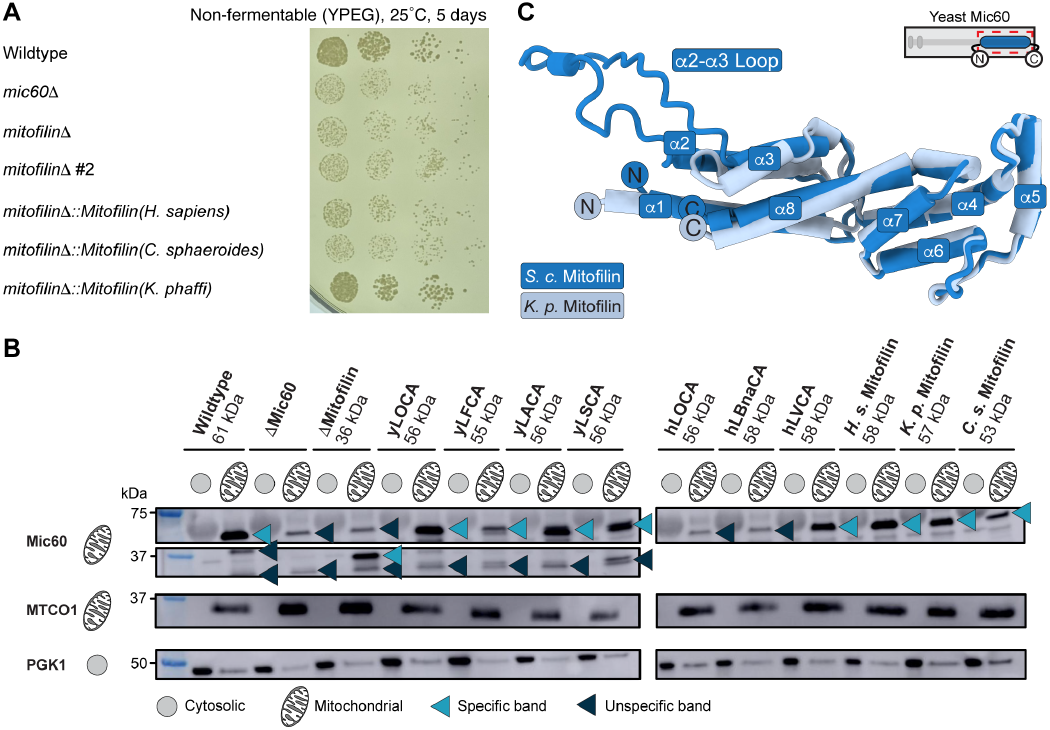
Respiratory growth phenotypes for yeast Mic60 Mitofilin chimeras. **(A)** Spot assay of yeast Mic60 chimeric strains on YPEG media, which is non-fermentable for yeast, after 5 days at 25°C. Deletion of the Mitofilin domain (*mitofilin*Δ and *mitofilin*Δ *#2*, which is the same strain as *mitofilin*Δ but a differerent colony) in yeast Mic60 or substituting it with the *H. sapiens* (*mitofilin*Δ*::Mitofilin(H. sapiens)*) and the *C. sphaeroides* (*mitofilin*Δ*::Mitofilin(C. sphaeroides*) Mitofilin domain results in knockout-like (*mic60*Δ) colony density and sizes. Yeast Mic60 with the *K. phaffi* Mitofilin domain (*mitofilin*Δ*::Mitofilin(K. phaffi*) exhibits wildtype-like growth. **(B)** Immunoblots of the cytosolic and the mitochondrial fractions of yeast Mic60 wildtype and chimeras after mitochondrial isolation against antibodies for Mic60 (top), cytochrome c oxidase subunit 1 (MTCO1, mitochondrial marker, center), and phosphoglycerate kinase (PGK1, cytosolic marker, bottom). All yeast chimeras except for hLOCA and hLBnaCA are expressed and localized to the mitochondria. Strains were grown for 28 hours in YPEG and the cell density was adjusted to an OD_600_ of 11 for each strain. The custom-made (Cusabio) polyclonal Mic60 antibody recognizes residues 112-325 of wildtype *S. cerevisiae* Mic60, thus extending 8 residues into the Mitofilin domain. Estimated molecular weights for the yeast Mic60 chimeras were obtained from the Expasy ProtParam Webserver^99,100^. **(C)** Superposition of AlphaFold2-predicted models of the *S. cerevisiae* Mitofilin domain (dark blue) and the *K. phaffi* Mitofilin domain (light blue) to highlight the extended Saccharomyceta clade loop region (‘α2-α3 Loop’) connecting helices α2 and α3.

Given that the structural conservation of the Mitofilin domain extends beyond all major eukaryotic groups into prokaryotic alphaproteobacteria (Figure 1B, C)^45^, we next asked whether Mitofilin-supported cellular respiration is similarly well conserved. We generated S288C strains where the *S. cerevisiae* Mitofilin sequence (residues 318-540) was swapped with that of Mitofilin homologues from different species at the endogenous locus. In contrast to previous reports^24^, we found that a chimeric yeast strain encoding the alphaproteobacterial *C. sphaeroides* Mitofilin domain (residues 270-433; *mitofilin*Δ*::Mitofilin(C. sphaeroides)*) shows a similar growth deficit to the *mic60*Δ and *mitofilin*Δ strains on non-fermentable media (Figure 2A). Similarly, the yeast encoding the human Mitofilin chimera (residues 562-758; *mitofilin*Δ*::Mitofilin(H. sapiens)*) displays a respiratory growth deficit like *mic60*Δ (Figure 2A). Western blots of isolated mitochondrial fractions confirm the mitochondrial localization of *mitofilin*Δ and the chimeric *H. sapiens* Mitofilin and *C. sphaeroides* Mitofilin yeast strains (Figures 2B). It is worth noting that a previous complementation study using plasmid-based expression observed a partial rescue of *S. cerevisiae* Mic60 function using the full-length *C. sphaeroides* Mic60 sequence^24^. In our work, we used CRISPR-Cas9 to generate chimeras of Mic60 at the endogenous locus of the *S. cerevisiae* genome. All strains were verified by whole genome sequencing (Figure S3).

### K. *phaffi* Mitofilin rescues respiratory growth in *S. cerevisiae*

We noticed in alignments of eukaryotic Mic60 sequences that the *S. cerevisiae* Mitofilin domain contains a loop region (residues 364-406) between a helices α2 and α3, which is distinct to the Saccharomyceta clade (Figure 2C). To test whether this loop is important for yeast Mitofilin respiratory function, we generated a *S. cerevisiae* strain that expresses a chimeric *S. cerevisiae* Mic60 with the Mitofilin domain of *K. phaffi* (residues 318-509; *mitofilin*Δ*::Mitofilin(K. phaffi)*), which lacks the unique loop region. We found that the Mic60 chimera with the *K. phaffi* Mitofilin domain localizes to mitochondria and grows like wildtype *S. cerevisiae* under non-fermentable conditions (Figure 2A, B). This indicates that despite only sharing 40% of sequence identity with *S. cerevisiae* Mitofilin (Figure 3B), the *K. phaffi* Mitofilin supports wildtype-like cellular respiration, and that the Saccharomyceta-specific loop region is not required for Mitofilin-supported respiratory growth in *S. cerevisiae*.

**Figure 3.**
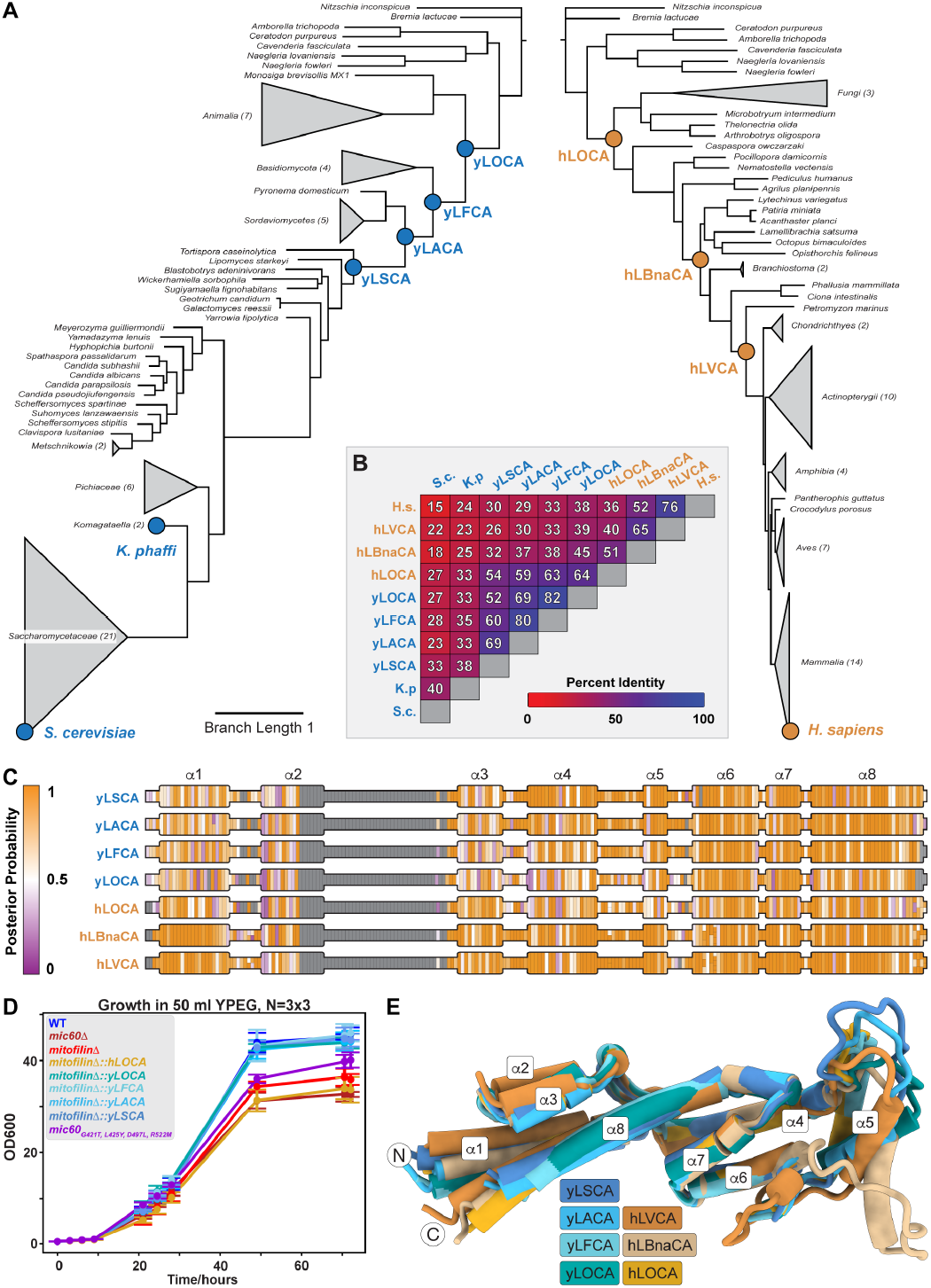
Ancestral sequence reconstruction of Mitofilin domains. **(A)** Yeast-focused (left) and animal-focused (right) phylogenies of the Mitofilin domain used for ancestral sequence reconstruction. Recon-structed nodes are designated by colored spheres (blue: yeast-derived ancestors; gold: animal-derived ancestors). Trees are scaled to identical branch lengths. Some clades are collapsed for clarity. Full trees with branch supports can be found in Figures S5 and S6. yLxCA: yeast-de-rived last common ancestor, hLxCA: human-derived last common an-cestor, LSCA: last Saccharomycotina common ancestor, LACA: last As-comycota common ancestor, LFCA: last Fungi common ancestor, LOCA: last ophisthokont common ancestor, LVCA: last vertebrate common ancestor, LBnaCA: last Bilateria non-arthropod common ancestor. Table showing pairwise sequence identity (in %) between each extant or reconstructed ancestral Mitofilin domain. Pairwise sequence identify is colored from red (low) to blue (high). **(C)** Linear topology schemes of reconstructed ancestors color-coded per residue by posterior probability. Posterior probability is colored from purple (low) to orange (high). Gray residues indicate absent residues in the reconstructed ancestor. Secondary structure elements (helices α1 to α8) are represented by protuberant regions. **(D)** Growth curve for wildtype yeast Mic60 (WT, blue), mic60Δ (dark red), mitofilinΔ (bright red), reconstructed Mitofilin ancestors (yellow, teal, and light blue colors), and mic60G421T, L425Y, D497L, R522M (purple) cultivated under respiratory conditions (50 ml YPEG; N = 9). **(E)** Superposition of AlphaFold2 models of reconstructed ancestors (dark blue: yLSCA; sky blue: yLACA; light blue: yLFCA, teal: yLOCA; gold: hLOCA; light brown: hLBnaCA; dark brown: hLVCA).

### Ancestral sequence reconstruction of Mitofilin ancestors

Our finding that *K. phaffi* Mitofilin rescues respiratory growth in *S. cerevisiae* suggests that the Mitofilin domain’s role in cellular respiration may be an ancient function, well-conserved in yeast. To test this hypothesis and to determine an evolutionary time point when yeast Mitofilin respiratory function diverged in opisthokonts, we used ancestral sequence reconstruction, a statistical phylogenetic approach,^43^ to infer the Mitofilin sequences of opisthokont ancestors. We then tested whether these Mic60 Mitofilin chimeras were sufficient to support respiratory growth.

We began by collecting Mic60 sequences from all eukaryotic clades, optimizing for phylogenetic coverage of Ascomycota fungi, and truncated the sequences to only contain the C-terminal Mitofilin domain as defined above. Following multiple sequence alignments, we applied a stringent amino acid substitution model to generate Mitofilin-based molecular phylogenies (Figure 3A, Figure S4). Briefly, substitution models were sorted by Bayesian information criteria^47^, and evaluated for their ability to achieve largely congruent evolutionary relationships between fungi (Figure S5). We found that the Q.yeast model^48,49^ best modeled this tree, with +I and +G4 decorators chosen to approximate the evolutionary rate^50^. Tanglegrams showed that mitofilin-based phylogenetic trees captured the structure of the Open Tree of Life, indicating that signal from the mitofilin sequences was sufficient to trace evolutionary trajectories (Figure S5). A yeast sequence-focused tree containing 121 sequences was then used to reconstruct ancestors along the fungal lineage from *S. cerevisiae* to the last common ancestors of Saccharomycotina (yLSCA), Ascomycota (yLACA), Fungi (yLFCA) and Opisthokonta (yLOCA) (Figure 3A-D). Reconstruction confidence is measured by the mean posterior probability for a reconstructed ancestor. We reconstructed 174-179 residues for yLSCA, yLACA, yLFCA and yLOCA Mitofilin with a mean posterior probability ranging from 72% for yLOCA to 82% for yLACA and yLSCA (Figure 3C, Figures S6).

We initially reconstructed one tree covering both yeast and animal sequences, but encountered long branch lengths, particularly for yeast Mitofilin sequences. This indicated that the available generalized substitution models did not accurately describe the evolutionary history of yeast Mic60. Therefore, we next reconstructed an additional tree with good phylogenetic coverage for animals to investigate the conservation of yeast respiratory function further down the lineage to the human Mitofilin domain (Figure 3A, Figures S5, S6, S7). Once more, tanglegrams for this human-lineage focused tree also captured relationships represented in the Open Tree of Life (Figure S5). This phylogenetic tree focused on human lineage-derived reconstructed Mitofilin domains for the last vertebrate (hLVCA), last Bilateria non-arthropod (hLBnaCA), and last opisthokont common ancestor (hLOCA) (see Methods for details). The mean posterior probability for hLVCA, hLBnaCA, and hLOCA ranged from 75-88% (Figure 3C, Figure S6). The yLOCA and hLOCA Mitofilin sequences share 64% sequence identify (Figure 3B).

### Yeast-lineage derived Mitofilin LOCA rescues respiratory growth

To evaluate the functional conservation of the ancestral Mitofilin sequences, we generated yeast strains encoding Mitofilin chimeras of Mic60, where we exchanged the wildtype Mitofilin domain with the reconstructed ancestral Mitofilin domains. All the reconstructed yeast-derived ancestors localize to the mitochondria, as shown by immunoblots of mitochondrial fractions (Figure 2B). Notably, all yeast ancestors (yLSCA, yLACA, yLFCA and yLOCA) grow like wildtype in respiratory media (Figure 3D, Figure S8). To better quantify yeast respiratory growth over time, we cultured yeast strains in liquid medium (50 ml YPEG) under non-fermentable conditions and assayed cell density by spectrophotometrically measuring the OD_600_ at defined time points. We observed two growth regimes: while growth rates do not significantly differ within the first 24 hours of growth, during mid-log phase, all yeast ancestors and *K. phaffi* Mitofilin grow like wildtype, whereas *mic60*Δ and *mitofilin*Δ show a significantly reduced growth rate, which is consistent with colony number and thickness in the solid growth assays (Figure 3D, Figure 2 for *K. phaffi*). Our findings suggest that Mitofilin’s respiratory growth function in yeast is conserved all the way to yLOCA, covering over one billion years of evolution^51^.

Whole genome sequencing confirmed the correct genomic insertion of the human-lineage derived ancestor sequences in the yeast strain; however, we did not observe stable expression for the human-derived hLBnaCA and hLOCA ancestors in immunoblots (Figure 2B). Accordingly, the human-lineage derived ancestors show deficient respiratory growth like *mic60*Δ (Figure 3D, Figure S8). While the hLVCA Mitofilin ancestor expresses and localizes to mitochondria (Figure 2B), it does not rescue respiratory growth (Figure S8).

### Structural comparison of ancestors reveals respiratory residue signature in yeast

The functional differences in our resurrected ancestral sequences allowed us to nominate common sequence elements to a conserved respiratory function. We first generated structure predictions of Mitofilin ancestors using our reconstructed sequences as input for AlphaFold2. As expected based on sequence identity, all predicted structures adopt the same overall fold (Figure 3E). To identify potential amino acids responsible for functional differences, we analyzed multiple sequence alignments of extant and resurrected Mitofilin sequences for similarities in functional constructs in yeast (wildtype, *K. phaffi* and all yeast-lineage derived ancestral sequences), compared with constructs that exhibited a KO-like growth phenotype (*H. sapiens* and all human-lineage derived ancestors) or did not stably express (hLOCA, hLBnaCA). We refer to functional constructs as extant and reconstructed sequences that show wildtype-like respiratory growth, and non-functional constructs as sequences that exhibit a *mic60*Δ-like respiratory growth curve or do not show stable expression in yeast (Figure 2B, Figure 3D, Figure S8).

We identified four residues in the Mitofilin domain, which are either identical or retain similar chemical properties (charge, hydrophobicity, size) in all functional constructs, but differ in the non-functional constructs (Figure 4A, B, Figure S9). Intriguingly, when mapped onto the predicted structure, these four residues create a line that runs across the center of the Mitofilin domain, perpendicular to the two distinctive four-helix bundles (Figure 4B). Two of these residues, Glycine 421 (*S. cerevisiae* nomenclature) and Leucine 425 are positioned on the loop between helices α3 and α4, which connects the predicted inner and outer 4-helix bundles of the Mitofilin domain (Figure 4B). Glycine 421 changes from a small (Glycine) or hydrophilic (Serine/Asparagine) residue in all functional sequences into a charged residue (Glutamate) in all human-derived ancestors. Notably, Glycine 421 corresponds to a Threonine (Threonine 622) in *H. sapiens*. Leucine 425 is a hydrophobic residue (Leucine/Alanine) in all yeast constructs, but a polar amino acid (Serine/Tyrosine) in all non-functional sequences. Aspartate 497 on a7 on the outer 4-helix bundle is a negatively charged or small polar residue (Aspartate/Glutamate/Serine) in functional constructs. In non-functional constructs, Aspartate 497 is a larger polar or a hydrophobic amino acid (Asparagine/Glutamine/Leucine). The fourth residue which differs between functional and non-functional residues,

**Figure 4.**
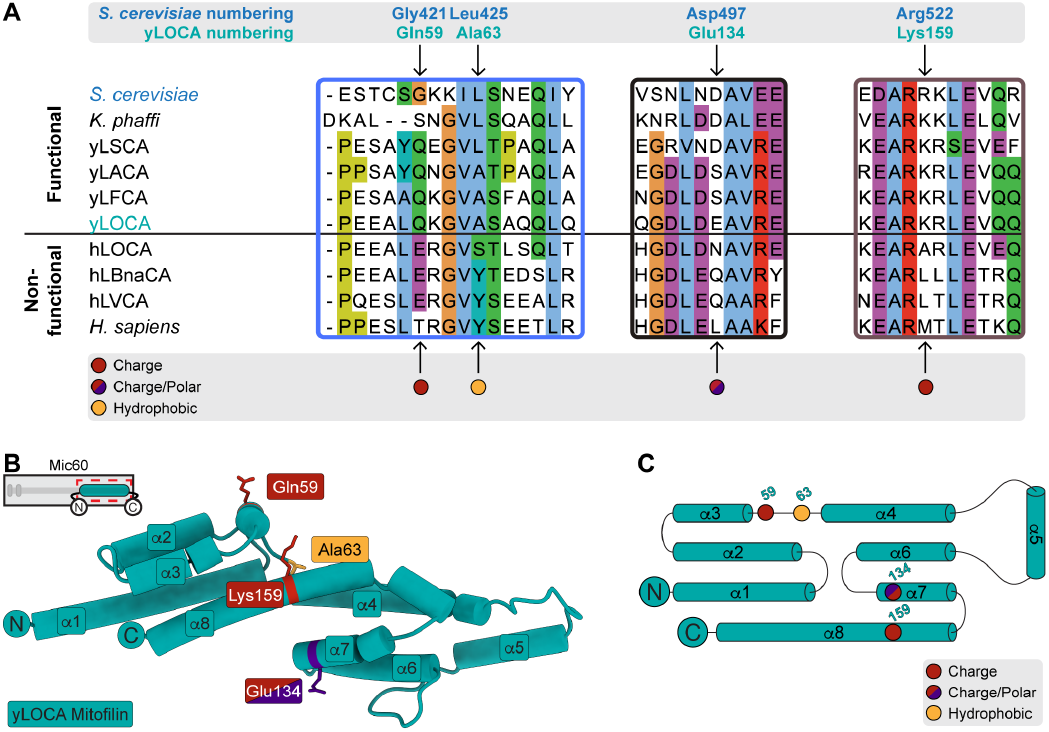
Four residues explain respiratory rescue in Mitofilin ancestors. **(A)** Contextualized regions of multiple sequence alignment between *S. cerevisiae, K. phaffi, H. sapiens*, and resurrected ancestor Mitofilin domains. Respiration functional and non-functional sequences are highlighted as such and separated by a horizontal black line. The four residues that explain respiratory growth phenotypes (G421, L425, D497, R522 in *S. cerevisiae* numbering) are indicated by arrows with *S. cerevisiae* and yLOCA numbering listed above the alignment. Chemical properties of the four ASR-identified residues in yeast and yeast-derived ancestors relative to human and animal-derived ancestors are denoted by colored spheres below the alignment (red sphere: change in charge; red/purple sphere: changes in charge and polar nature; yellow sphere: change in hydrophobicity). Colored boxes in alignment correspond to colored boxes in full alignment in Figure S9. **(B)** The ASR-identified residues are mapped onto the AlphaFold2 predicted yLOCA model. Residues are numbered by yLOCA nomenclature (see (A) for conversion to *S. cerevisiae* numbering) and colored by their chemical properties as described in (A). All four residues map to the central area of the Mitofilin domain. **(C)** Topology diagram of yLOCA showing the location of the four ASR-identified residues, which are colored by their chemical properties.

Arginine 522, is located on helix α8 and exists as a positively charged residue in functional sequences (Arginine/Lysine), and a hydrophobic residue (Alanine/Leucine/Methionine) in non-functional constructs.

### A four-residue signature is important for respiratory growth in yeast

To test whether the four residues identified by ancestral sequence reconstruction impact respiratory growth, we mutated these four residues in wildtype yeast Mic60 to their non-functional animal counterpart in the endogenous Mic60 gene (*mic60*_*G421T, L425Y, D497L, R522M*_). By measuring respiratory growth in liquid medium under non-fermentable conditions (50 ml YPEG), we observed that mutating just the four candidate residues in the otherwise unmodified wildtype yeast Mic60 results in impaired respiratory growth (Figure 3D), confirming that the residue signature, consisting of G421, L425, D487, and R522M, is important for Mic60’s role in mitochondrial respiration.

### Mitochondrial ultrastructural and network phenotypes of ancestors and mutants

Several TEM studies have shown that Mic60 deletion, as well as truncation of the Mitofilin domain, results in detached cristae and the loss of crista junctions in mitochondria^11,12,18,42,52^. Consistent with previous reports, we observed mitochondria without a clear cristae junction attachment in the *mic60*Δ and *mitofilin*Δ strains (Figure 5A). Surprisingly, however, the majority of mitochondria of functional, yeast-derived Mitofilin ancestor strains (yLOCA), which rescue respiratory growth like wildtype (Figure 3D), feature knockout-like detached cristae under both fermentative and respiratory growth conditions (Figure 5A, Figure S10).

**Figure 5.**
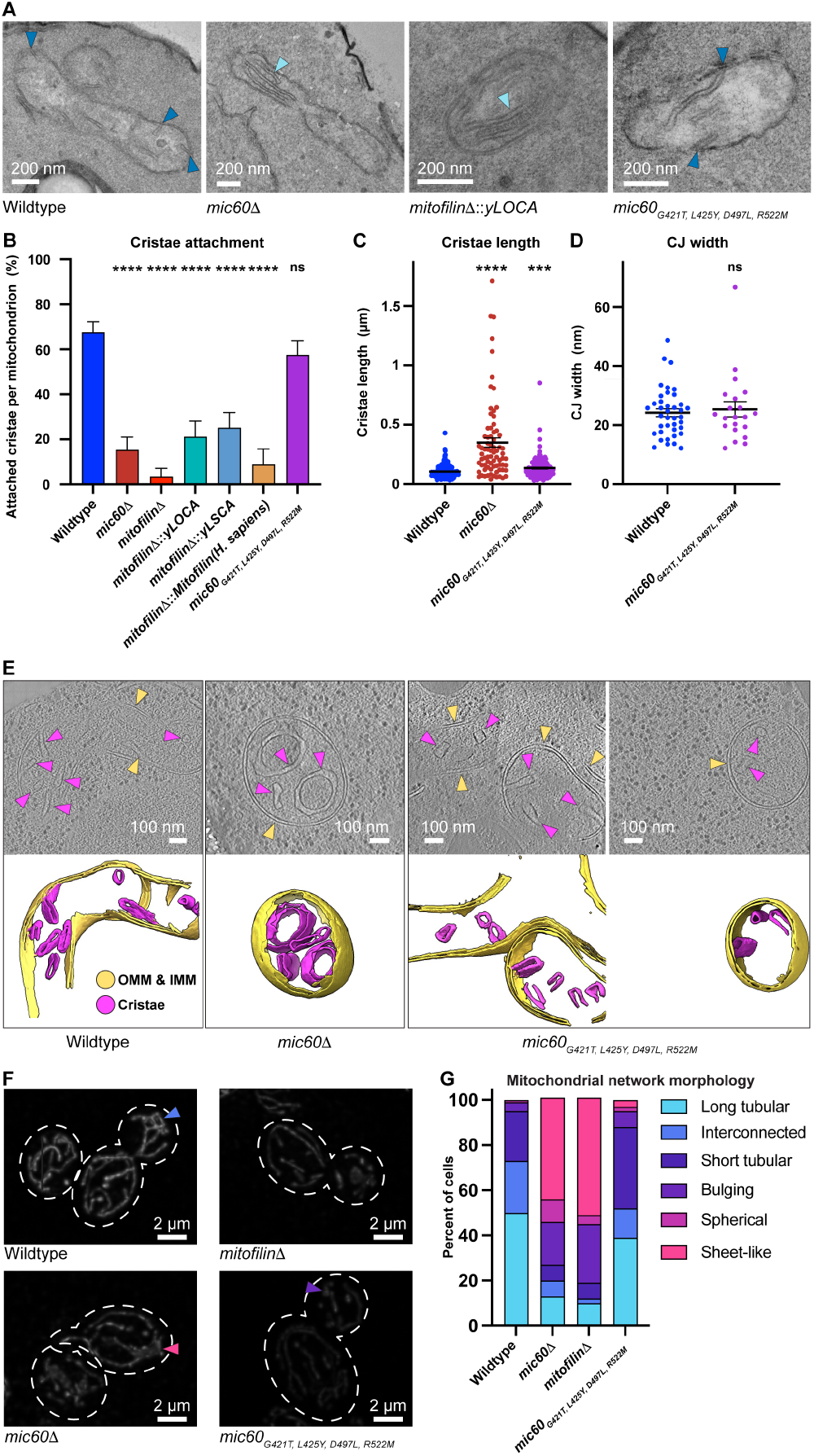
Transmission electron micrographs suggest lack of correlation between respiratory function and cristae attachment. **(A)** Representative TEM micrographs of mitochondria in WT *S. cerevisiae* (top left), *mic60*Δ (bottom left), *mitofilin*Δ*::yLOCA* (top right), *mic60*_*G421T, L425Y, D497L, R522M*_ (bottom right). Scale bar: 200 nm. **(B)** Percentage of attached cristae per mitochondrion. N equals the total number of mitochondria counted in separate cells: Wildtype = 43, *mic60*Δ = 23, *mitofilin*Δ = 14, *mitofilin*Δ*::yLOCA* = 22, *mitofilin*Δ*::yLSCA* = 33, *mitofilin*Δ::*Mitofilin(H. sapiens)* = 15, *mic60*_*G421T, L425Y, D497L, R522M*_ = 38. Significance of difference is tested relative to wildtype using Welch’s t test; ****p < 0.0001. **(C)** Quantification of cristae length in indicated yeast strains shown in (A). Numbers of measured cristae in mitochondria from separate cells are represented as scatter plots with mean ± SEM. N refers to the number of cristae: Wildtype = 212, *mic60*Δ = 75, *mic60*_*G421T, L425Y, D497L, R522M*_ = 137. Significance of difference is tested relative to wildtype using Welch’s t test; ****p < 0.0001; ***p = 0.0008. **(D)** Quantification of CJ width in indicated yeast strains shown in (A). Values of CJ width measured in mitochondria from separate cells are shown as scatter plots with mean ± SEM. N refers to the number of CJ measured: Wildtype = 39, *mic60*_*G421T, L425Y, D497L, R522M*_ = 21. Significance of difference is tested relative to wildtype using Welch’s t test; p = 0.6876. **(E)** (Top) Representative summed projections of central slices of cryo-electro tomograms of mitochondria in strains indicated. (Bottom) 3D segmentations of mitochondrial membranes in the indicated yeast strains. **(F)** Representative maximum projections of mitochondrial morphology in indicated *S. cerevisiae* strains labeled with PKmito Orange. Triangles denote examples of different morphotypes: inter-connected tubules (blue), sheet-like (pink), bulbous (purple). **(G)** Quantification of mitochondrial morphology of cells shown in (F). A minimum of 90 cells were analyzed for each strain and data are represented as mean percentage of cells.

For the respiration-impaired 4-residue mutant *mic60*_*G421T, L425Y, D497L, R522M*_, TEM showed that while the number of attached cristae and the mean cristae junction width is similar to that of wildtype, the cristae of *mic60*_*G421T, L425Y, D497L, R522M*_ are longer than those of wildtype (Figure 5A-D). In tomograms of vitrified mitochondria, we observed that several cristae of *mic60*_*G421T, L425Y, D497L, R522M*_ show box-like shapes with sharp corners, a phenotype not observed in wildtype and the knockout strains (Figure 5E). Finally, the gross mitochondrial network morphology of *mic60*_*G421T, L425Y, D497L, R522M*_ resembles that of wildtype, with the majority of mitochondria forming a tubular network consisting of long, short, and interconnected mitochondria (95% of wildtype cells, 88% of *mic60*_*G421T, L425Y, D497L, R522M*_ cells; Figure 5F, G). This is in contrast to the knockout strains, of which a significant number of cells contain mitochondria with a sheet-like phenotype, as shown previously for *mic60*Δ (45% of *mic60*Δ cells, 52% of *mitofilin*Δ cells; Figure 5F, G)^26^.

Together, our observations reflect a lack of correlation between respiratory function of Mic60 ancestor and mutant strains and cristae attachment as assessed by TEM. Instead, we observe other cristae morphological defects, such as longer average cristae length and altered cristae shape associated with compromised respiratory function.

## Discussion

Respiration is a key function supported by mitochondrial cristae. While the requirement of Mic60 for stable cristae junctions is well-established^27,29,42^, understanding the sequence determinants for cristae-supported respiration has been lacking. Here we used ancestral sequence reconstruction to infer Mic60 Mitofilin sequences along the fungal lineage from *S. cerevisiae* to the last opisthokont common ancestor (LOCA). We show that Mic60 Mitofilin ancestors as far back as the last opisthokont common ancestor (yLOCA) rescue yeast mitochondrial respiration. This observation is striking, given the high rate of Mitofilin evolution in fungi, as indicated by the number of residue substitutions in the yeast phylogenetic tree compared to that of animals.

Previously, Tarasenko *et al*. reported a “slight” rescue of yeast respiratory growth by plasmid-expressed and mitochondrially targeted wildtype C-terminally FLAG-tagged full-length alphaproteobacterial Mic60^24^. In contrast, when introducing the Mitofilin domain of the same alphaproteobacterial species as a yeast chimera at the endogenous locus with CRISPR-Cas9, we observed a clear *mic60*Δ-like respiratory growth deficit. Since Mic60 function is strongly influenced by expression levels^30,32^, we opted for genome editing by CRISPR-Cas9 at the endogenous locus as opposed to plasmid-based expression. In addition, despite confirmation of correct insertion by whole-genome sequencing, two yeast Mic60 chimeras with human-lineage derived Mitofilin ancestors do not show protein expression (Figure 2B). Accordingly, these Mic60 chimeras also display a *mic60*Δ-like respiratory growth phenotype (Figure 3D, Figure S8). Since the expressed and non-expressed yeast Mic60 chimeras all contain the N-terminal mitochondrial targeting sequence and only differ in the C-terminal Mitofilin domain (Figure 1A), these observations suggest that additional, yet to be defined, mechanisms of proteostasis may be at play.

To more accurately trace the evolutionary history of Mic60, we generated two phylogenetic trees–one reflecting the evolutionary trajectory of yeast Mic60 sequences, and the other one focusing on the evolution of animal Mic60 sequences. We noticed that in a single tree of opisthokont Mic60 sequences, Mic60 evolution (in particular, the rapidly evolving yeast lineages) was insufficiently described by the available generalized amino acid substitution models. Furthermore, given that the Bayesian information criterion (BIC) used to accurately assess the quality of the substitution model requires a low number of sequences to meet statistical conditions, generating two trees allowed us to increase the number of sequences and optimize species representation^53^. As a probabilistic approach, ancestral sequence reconstruction includes an inherent degree of uncertainty, even with the most robust reconstructions. By correlating respiratory growth profiles with the corresponding amino acid sequences, we were able to identify just four residues that explain the difference in Mitofilin function for yeast-and animal-derived ancestors. This precision is an advance over previous structure-function dissections of the Mitofilin domain, which compared truncation constructs^27,29,42^. We confirmed the respiratory impact of the 4-residue signature in respiratory growth assays. The chemical properties of these residues differ in functional and non-functional constructs. Interestingly, these residues are also predicted to be close to each other in the tertiary structure, bisecting the central axis of the Mitofilin domain, perpendicular to the two distinctive four-helix bundles.

While the four residues do not impact CJ number and width, they affect cristae growth, resulting in slightly longer cristae than the wildtype, as assessed by TEM. In addition, we note a difference in cristae morphology as visualized by cryo-ET. Intriguingly, the decoupling between respiratory growth and the wildtype-like cristae junction attachment and morphology has been observed in other contexts, such as in *T. brucei*^54^. We note that while our TEM analysis was done under fermentative and respiratory conditions, our mitochondrial network observations were made under non-respiratory conditions, as done by others^26^. Future investigation under different conditions and with more sensitive methods will be necessary to fully understand Mic60-driven cristae biogenesis.

A potential source for compromised respiratory function in *mic60*_*G421T, L425Y, D497L, R522M*_ may be disrupted protein-protein interactions. These residues are adjacent to the previously described Mic19 binding site (Figure S11). Previous work using chemical cross-linking mass spectrometry suggest that in mice, Mic10 and Mic13 bind in the vicinity of the four residues^33^. From comparisons between the mitochondrial inner membrane fusogens Mgm1 in yeast and its animal ortholog Opa1, we note that differences in binding partners may be an important consideration. While these dynamin family proteins may show structural similarity, divergent interaction interfaces may result in variable epistasis, and distinctive inner membrane remodeling mechanisms^55^.

Given Mic60’s functions, candidate roles for the four residues we identified may include self-assembly, or protein-protein interactions. While Mic60’s Mitofilin domain interacts with several proteins, its binding interfaces remain obscure^33-35^. By identifying functionally important residues, this study lays the foundation for future work investigating mechanistic details of Mic60-supported respiration.

## Acknowledgements

We thank L. Stirling Churchman for gifting us the S288c DBY12045 yeast strain; Heike Rampelt and Jan Brix for gifting us Mic19 and Mic60 antisera; Ulandt Kim and the NextGen Sequencing Core at MGH for their outstanding support; Andrew J. Roger and Sergio A. Muñoz-Gómez for helpful discussions and advice. We thank Paula Montero Llopis and the Microscopy Resources on the North Quad (MIcRoN) Core at Harvard Medical School for support in fluorescent imaging. We are grateful to Sarah Sterling and Jennifer Podgorski at the MIT.nano cryo-EM facility for assistance in training and data collection. We acknowledge support from the Swiss National Science Foundation grant P180777 (F.M.C.B.), the Keystone Future of Science Fund (F.M.C.B), the Helen Hay Whitney Foundation (T.A.B.), Natural Sciences and Engineering Research Council of Canada grant RGPIN-2016-04801 (C.J.B.d.C.), Canada Foundation for Innovation grant 34475 (C.J.B.d.C.), Canadian Institutes of Health Research grant 377068 (C.J.B.d.C.), New Frontiers in Research Fund-Exploration grant NFRFE-2018-00064 (C.J.B.d.C.), the Moore–Simons Project on the Origin of the Eukaryotic Cell grant 9736 (L.H.C., C.J.B.d.C.) and National Institutes of Health grant R35GM142553 (L.H.C.).

## Author contributions

F.M.C.B., T.A.B., T.H.N., D.S. and L.H.C. conceived of the study and designed the experiments. F.M.C.B., T.A.B., T.H.N. and D.S. generated the yeast strains. F.M.C.B. and T.H.N. performed growth assays and mitochondrial localization experiments. T.A.B., T.H.N, L.B.C and C.J.B.d.C. performed ancestral sequence reconstruction. M.C., A.E.H.N. and M.E. performed the TEM consultation and imaging. L.H.C prepared the cryo-ET samples and collected the data. Y.-T.L. processed the cryo-ET data. M.P.K. contributed to mutant analysis. F.M.C.B., T.A.B., T.H.N., C.J.B.d.C., and L.H.C. analyzed the data. F.M.C.B. and T.A.B. created figures and tables, and F.M.C.B. and L.H.C. wrote the manuscript with contributions from T.A.B. and T.H.N. All authors approved the final version of the manuscript.

## Competing interest statement

The authors declare no competing interests.

## Materials and Methods

### Yeast Strains and Growth

All *S. cerevisiae* strains used in this work are derivatives of the S288c DBY12045^56^ strain (kindly gifted by L. Stirling Churchman). A complete list of all yeast strains generated and used is included as Table S2.

For spot assays, yeast strains were first grown in YPD (1% yeast extract, 2% peptone, 2% glucose) or YPEG (1% yeast extract, 2% peptone, 2% ethanol, 3% glycerol) media at 30°C to an OD_600_ of 0.2 to 0.3. Cells were pelleted by centrifugation at 6,000 x g for 2 min and resuspended in water to a final OD_600_ of 0.1. Each sample was 5-fold serially diluted in water to obtain cells with an OD_600_ value ranging from 0.1 to 6.4 × 10^-6^, then transferred to YPEG or YPD agar plates using a multichannel pipette. Plates were incubated at 30°C, 25°C, or 18°C and imaged at 24-hour intervals for 7 days.

For liquid growth assays, yeast strains were first grown in YPD at 30°C to an OD_600_ of 11. Following centrifugation at 6,000 x g for 2 minutes, cell pellets were resuspended in 50 ml YPEG to a final OD_600_ of 0.4 and grown at 30°C for 72 hours. 3 technical and 3 biological replicates (N=9) were grown for each strain. Measurements were taken at defined time points and analyzed using the matplotlib^57^ and pandas^58^ libraries in Python.

### CRISPR/Cas9 Endogenous Genome Editing

Mic60 knockout (*mic60*Δ), Mitofilin knockout (*mitofilin*Δ), chimeric yeast Mic60 strains expressing Mitofilin domains of other species or reconstructed ancestral Mitofilin domains, and *mic60*_*G421T, L425Y, D497L, R522M*_ were generated by CRISPR-Cas9 using the method by Levi and Arava^59^. Synthesized oligonucleotides (Integrated DNA Technologies) encoding the desired sgRNA sequence were cloned into the Cas9-encoding bRA66 plasmid (Addgene, 100952). Yeast strains were co-transformed with 500 ng of modified bRA66 plasmid (encoding Cas9 and sgRNA) and 5 ug of synthesized donor DNA (GenScript) using a lithium acetate/PEG-3350/heat-shock protocol described by Shaw *et al*^60^. All plasmids, sgRNAs and donor DNAs are listed in Table S3. Gene modifications were verified by sequencing.

### Whole Genome Sequencing Analysis

Genomic DNA from each yeast strain was extracted using the MasterPure Yeast DNA Purification Kit (Biosearch Technologies) and sequenced with HiSeq 50 nt paired-end reads (Illumina). Reads were aligned against the *S. cerevisiae* S288C genome (Genbank GCF_000146045.2) using bwa^61^ and samtools/bcftools^62^. For chimeric constructs, custom reference genomes were produced by manually editing the *S. cerevisiae* S288C genome in fasta format. Aligned reads were visualized using Integrated Genome Browser^63^. Chromosomal coverage was assessed with custom python scripts, available in an open-source repository (https://github.com/tribell4310/wgs_pipe).

### Immunoblot Analysis of Protein Localization

Expression and mitochondrial localization of Mic60 for all yeast strains were evaluated through antibody detection by immunoblotting of isolated mitochondria. Yeast strains were prepared and grown as described above for growth assays in liquid YPEG for 28 hours. After centrifugation at 1,500 x g for 5 minutes, mitochondria were isolated using the Yeast Mitochondria Isolation Kit (Abcam). Spheroblasts were homogenized by pushing 12 times through a 25G needle. For immunoblot analysis, the following antibodies were used: anti-Mic60 (custom-made, Cusabio), Mic60 antisera, Mic19 antisera (both antisera were kindly gifted by Heike Rampelt and Jan Brix), anti-MTCO1 as a mitochondrial marker (ab110270, Abcam) and anti-PGK1 as a cytosolic marker (ab113687, Abcam).

### Ancestral Sequence Reconstruction

Ancestral reconstruction was carried out following the method of Prinston *et al*.^64^. Briefly, Mic60 sequences were curated from the InterPro database^65^ (IPR019133) as well as through searches of NCBI GenBank^66^ and Uniprot^67^ using BLAST^68^ and HMMER^69^ algorithms. Searches returned >5,000 sequences, which were aligned with MAFFT^70^. Sequences with large insertions or deletions and sequences with greater than 96% pairwise sequence identity were eliminated immediately. This set of sequences (Table S4) was then whittled down to 69 (yeast-focused tree) or 62 (animal-focused tree) sequences that effectively represented the full phylogenetic diversity of the evolutionary space using custom python scripts, available in an open-source repository (https://github.com/tribell4310/phylogenetics). Seven amoebozoan and other non-opisthokont eukaryotic sequences were then added to the alignments for outgroup rooting, and the sequences were re-aligned using PRANK^71^. The best-fit substitution model (Q.yeast^48,49^ with allowed invariable sites (+I) and a discrete Gamma model of site heterogeneity^72^ with four rate categories (+G4) for the yeast-focused tree; LG^73^ with a four-class free-rate model^50,74^ of site heterogeneity (+R4) for the animal-focused tree) was determined using MODELFINDER^75^ according to Bayesian information criterion. Phylogeny construction and ancestral reconstruction were performed in IQ-TREE^76^ using default settings. Branch supports were inferred using SH-like approximate likelihood ratio tests^77^ in IQ-TREE. Tanglegram comparison of phylogenies with the Open Tree of Life^78^ was performed using the rotl package^79^, as well as custom R scripts made available in the above-referenced open-source repository. All alignment positions were reconstructed without regard to relatively unpopulated gap regions. Gaps were manually removed from the reconstructed sequences by comparing the reconstructions to their nearest-neighbor extant sequences using a parsimony approach similar in principle to Fitch’s algorithm^80^.

### Structure Modeling and Analysis

Structure predictions for all proteins shown were generated by AlphaFold2^46^ using ColabFold v. 1.5.5^81^. All sequences used for protein predictions are listed in Table S1. Visual representations of protein structures were created and analyzed with UCSF ChimeraX (Resource for Biocomputing, Visualization, and Informatics, University of California, San Francisco)^82^ and PyMOL (The PyMOL Molecular Graphics System, Version 2.5 Schrödinger, LLC). Protein folds were compared by superposition using the matchmaker^83^ command in ChimeraX. Sequence conservation was mapped onto the structures using the ConSurf webserver^84,85^.

### Transmission electron microscopy

Cells for TEM imaging were prepared according to an adaptation of the method by Bauer *et al*^86^. Yeast strains were cultured in either YPD or YPEG medium (respiratory conditions are indicated in each figure legend) for 16 hours at 30°C, harvested by centrifugation at 6,000 x g for 30 seconds, and washed in cold phosphate-buffered saline (PBS, pH 7.2). The cells were then fixed for 30 minutes with 2% glutaraldehyde in 0.1 M sodium cacodylate buffer (pH 7.2) at 4°C. Following three washes with 0.1 M cold sodium cacodylate buffer, the cells were incubated in 50 mM Tris-HCl at pH 7.5, 5 mM magnesium chloride, 1.4 M sorbitol, 0.5% (v/v) 2-mercaptoethanol, and 0.15 mg/ml Zymolase 20T for 10 minutes at room temperature for digestion of the cell wall. After four washes with 0.1 M sodium cacodylate buffer, the cell pellets were post-fixed in 1% Osmiumtetroxide (OsO_4_)/1.5% Potasiumferrocyanide (KFeCN_6_) for 1 hour, washed 2x in water, 1x Maleate buffer (MB) and incubated in 1% uranyl acetate in MB for 1 hour. Pellets were then rinsed in ddH_2_O and dehydrated through a series of ethanol (50%, 70%, 95%, (2x)100%) for 15 minutes per solution. Dehydrated samples were put in propylene oxide for 5 minutes before they were infiltrated in Spurr resin mixed 1:2 with propylene oxide overnight at 4°C, followed by 100% Spurr resin overnight at 4°C. Samples were polymerized in a 60°C oven in Spurr resin for 48 hours. They were then sectioned into 80 nm thin sections, sections were picked up on formvar/carbon coated grids, stained with 0.2% lead citrate, and imaged on a TecnaiBioTwin Transmission Electron Microscope equipped with an AMT Nanosprint 43-MKII camera.

#### Quantitative analysis

Attached and detached cristae were counted manually on TEM micrographs. Cristae length and CJ width were measured on TEM micrographs using the line selection tool in FIJI^87^. Statistical analyses were carried out using the Prism 10 GraphPad software.

### Fluorescence microscopy

Yeast strains were cultured in YPD medium for 16 hours at 30°C, harvested by centrifugation at 6,000 x g for 30 seconds, and stained with 1:1000 PKmito Orange (Spirochrome) by incubation at 37°C for 30 minutes. Prior to imaging, PKmito Orange was replaced with YPD media. Images of live mitochondria on 2% YPD-agarose pads were acquired on a fully automated spinning disk confocal microscope, consisting of a Yokogawa CSU-W1 scan unit mounted on a fully automated Nikon Ti2 inverted stand, enclosed within an OKOLab environmental chamber set to 37°C and equipped with an Andor Zyla 4.2Plus sCMOS camera. Z-stacks were acquired using a Plan Apo 100x/1.45 oil immersion (Cargille 37 immersion media) using a MadCity Labs Piezo stage insert. Z-step was set to 200 nm, and the total range was ∼6.5 `µ`m. Signal from pKMito was obtained by exciting the fluorophore with a DPSS 561 nm laser (Nikon LUN-F combiner; 100 mW at the fiber), using a Semrock Di01-T405/488/568/647 dichroic mirror and a Chroma ET605/52m emission filter. Images were collected at 16 bit dual gain. Nikon Elements AR Acquisition Software (version 5.41.02) was used to acquire the z-stacks.

In addition to the mitochondria, z-stacks of carboxylate red (580/605nm) 100 nm fluorescent beads were acquired to measure the point spread function (PSF) of the system. These PSF stacks were used to deconvolve the pKMito z-stack images using Huygens deconvolution software, outfitted with the spinning disk module.

#### Quantitative analysis

Mitochondrial network morphology was assessed in FIJI^87^ using deconvoluted z-stacks by manually sorting cells into six morphological categories based on previous classifications used by Friedman et al^26^: “Long tubular” describes cells with predominantly elongated mitochondria, which are not fused into a mesh. “Short tubular” refers to cells containing shorter mitochondria. “Interconnected” depicts cells that contain elongated mitochondria with several branching points forming an interconnected mesh. “Bulging” describes cells containing mitochondria with bulbous sections. “Spherical” refers to cells containing a mixture of fragmented-looking round mitochondria and tubular mitochondria. “Sheet-like” is used for cells with several flat, wide mitochondria.

### Cryo-electron tomography

#### Specimen preparation

Yeast cells were cultured in YPD medium for 16 hours at 30°C. 3 `µ`l of yeast cells diluted to an OD_600_ of 1.5 were deposited onto Quantifoil Cu R 1.2/1.3 grids (Electron Microscopy Sciences) and glow discharged for 90 seconds at 15 mA. Grids were back-blotted for 3 seconds after a 5 second wait time using a FEI Vitrobot Mark IV (Thermo Fisher Scientific) at room temperature, 100% humidity, blotting force of 15, and then plunged into liquid ethane. Subsequent grid handling and transfers were performed in liquid nitrogen. Grids were clipped into cryo-FIB autogrids (Thermo Fisher Scientific).

#### Cryo-FIB milling

Clipped grids were loaded into an Aquilos 2 (Thermo Fisher Scientific), sputter coated with inorganic platinum, then GIS coated in organometallic platinum. Cryo-FIB milling was performed to generate two rectangular patterns to mill top and bottom sections of cells, with adjacent two micro-expansion joints to improve lamellae stability^88^. Cryo-FIB milling was conducted at a nominal tilt angle of 12-18°, which translates to a milling angle of 7-10° and in steps of decreasing ion beam current (1 nA = 10 pA) and thickness to achieve 150-250 nm lamellae.

#### Data collection

Data were collected using the automated data collection software SerialEM^89^ on a FEI Titan Krios equipped at 300keV with an energy filter and Gatan K3 direct detector. Images were collected with a defocus range of 4-6 `µ`m using super resolution mode with a calibrated pixel size of 1.375 Å/pixel. Tilt range was -60 to 60° in 2° increments following a Hagen dose-symmetric scheme^90^, with a total dose of 141 electrons/ Å^2^.

#### Cryo-ET data processing, tomogram reconstruction and segmentation

Data were preprocessed using TOMOgram MANager (TOMOMAN) wrapper scripts^91^, including MotionCor2^92^, dose filtering, and CTFFIND4^93^. Alignment and reconstruction were then performed with AreTomo (v1.3.0)^94^. The bin4 (11 Å/pixel) tomograms were denoised using cryo-CARE^95^ and missing-wedge corrected with IsoNET^96^. Tomogram segmentation was carried out using Membrain-Seg^97^ (pretrained model v9). All segmentations were subsequently manually refined in Amira 2023.2 (Thermo Fisher Scientific) and then imported into ChimeraX^82^ 1.8 using the ArtiaX^98^ plugin for visualization and figure generation.

## Notes

### Competing Interest Statement

The authors have declared no competing interest.

### Summary of Updates

Revision 1: Addition imaging experiments (Figure 5), updated Figure 2.

