## Supplement for "Ancestral sequence reconstruction of the Mic60 Mitofilin domain reveals residues supporting respiration in yeast"

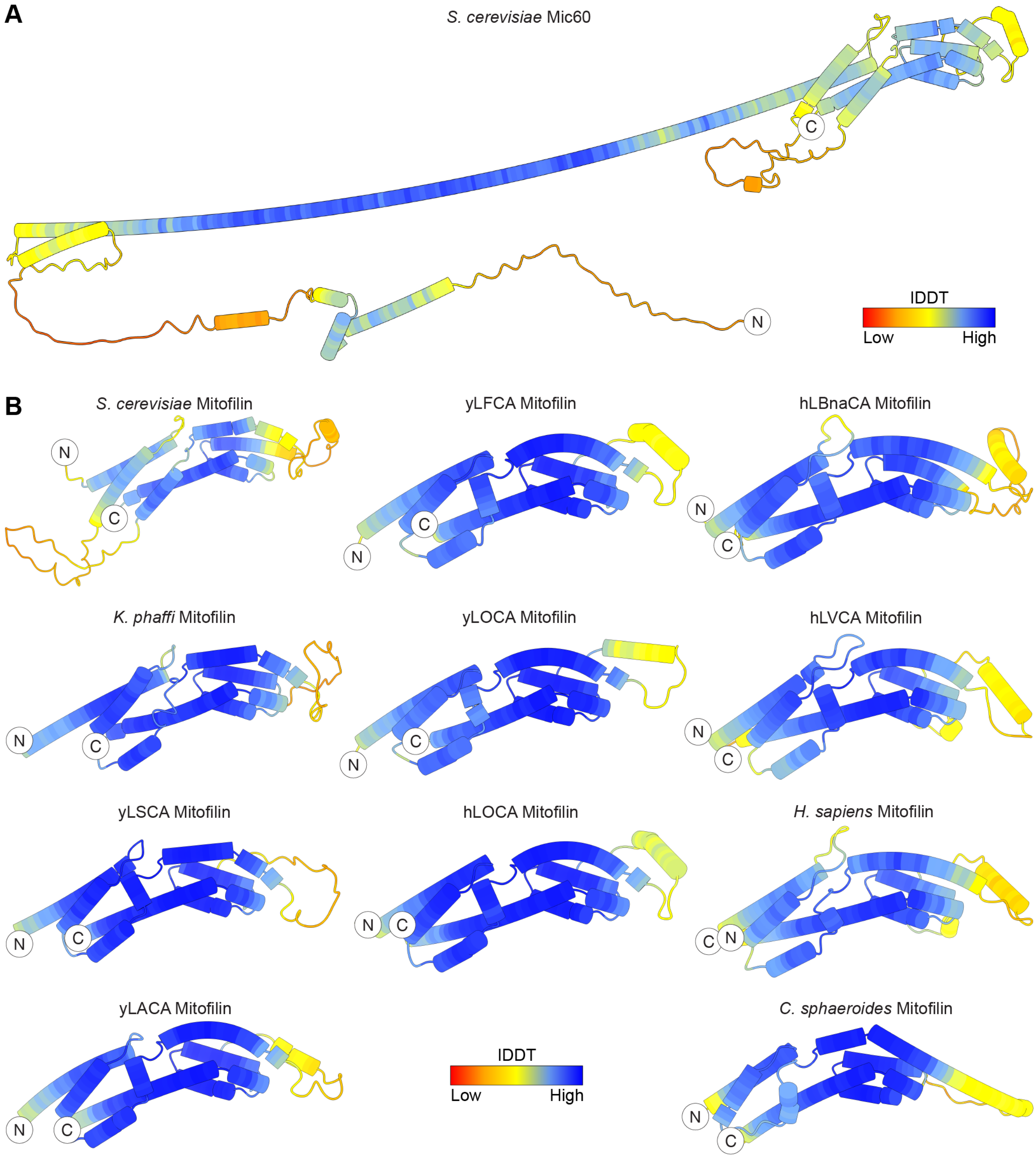


**Figure S1: The title of a supplementary figure. (A)** (Structure prediction of *S. cerevisiae* Mic60 (AF-P36112-F1)[1,2] colored by prediction confidence from low (red) to high (blue). Prediction confidence is expressed by a color-assigned IDDT-score[3]. **(B)** Mitofilin structure predictions colored by prediction confidence from low (red) to high (blue)[1,4]. Abbreviations: yLxCA: yeast-derived last common ancestor, hLxCA: human-derived last common ancestor, LSCA: last Saccharomycotina common ancestor, LACA: last Ascomycota common ancestor, LFCA: last Fungi common ancestor, LOCA: last ophisthokont common ancestor, LVCA: last vertebrate common ancestor, LBnaCA: last Bilateria non-arthropod common ancestor.

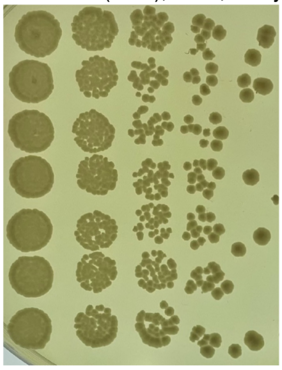


**Figure S2. Mitochondria-independent growth on fermentable media is not impacted by yeast Mic60 mutations.** Spot assay of yeast Mic60 strains on fermentable YPD agar (1% yeast extract, 2% peptone, 2% glucose) after 5 days at 25˚C, showing that Mic60 mutations do not impact mitochondria-independent cellular growth. *mic60Δ*: Mic60 knockout, *mitofilinΔ*: Mic60$_{1-317}$, *mitofilinΔ #2:* same strain as *mitofilinΔ* but different colony, *mitofilinΔ::Mitofilin(H. sapiens)*: yeast Mic60$_{1-317}$ with the human Mic60 Mitofilin sequence (residues 562-758 of *H. sapiens* Mic60), *mitofilinΔ::Mitofilin(C. sphaeroides)*: yeast Mic60$_{1-317}$ with an alphaproteobacterial Mic60 Mitofilin sequence (residues 270-433 of *C. sphaeroides* Mic60), *mitofilinΔ::Mitofilin(K. phaffi)*:: yeast Mic60$_{1-317}$ with the *K. phaffi* Mic60 Mitofilin sequence (residues 318-509).

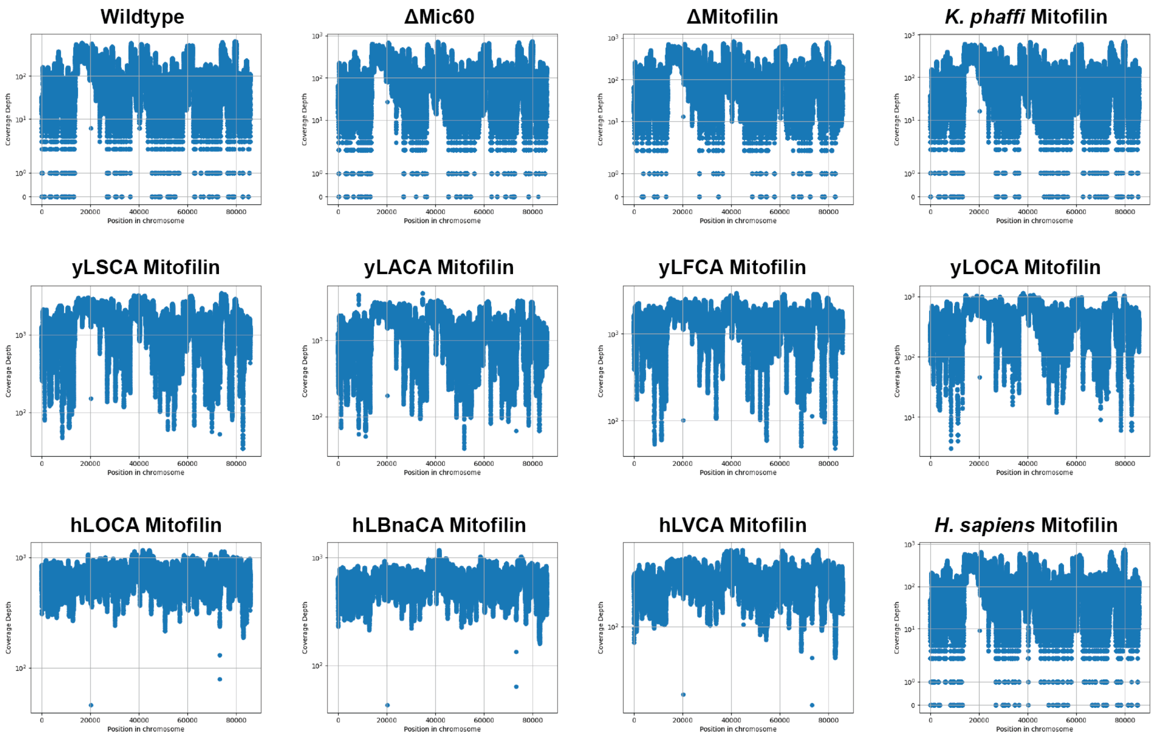


**Figure S3. Whole genome sequencing of yeast Mic60 chimeras reveals presence of mitochondrial DNA in all strains.** Plots of sequencing coverage across the mitochondrial chromosome for each yeast Mic60 strain used in this study. None of the respiratory defects observed in these strains are attributable to loss of mitochondrial DNA (i.e., the petite phenotype).

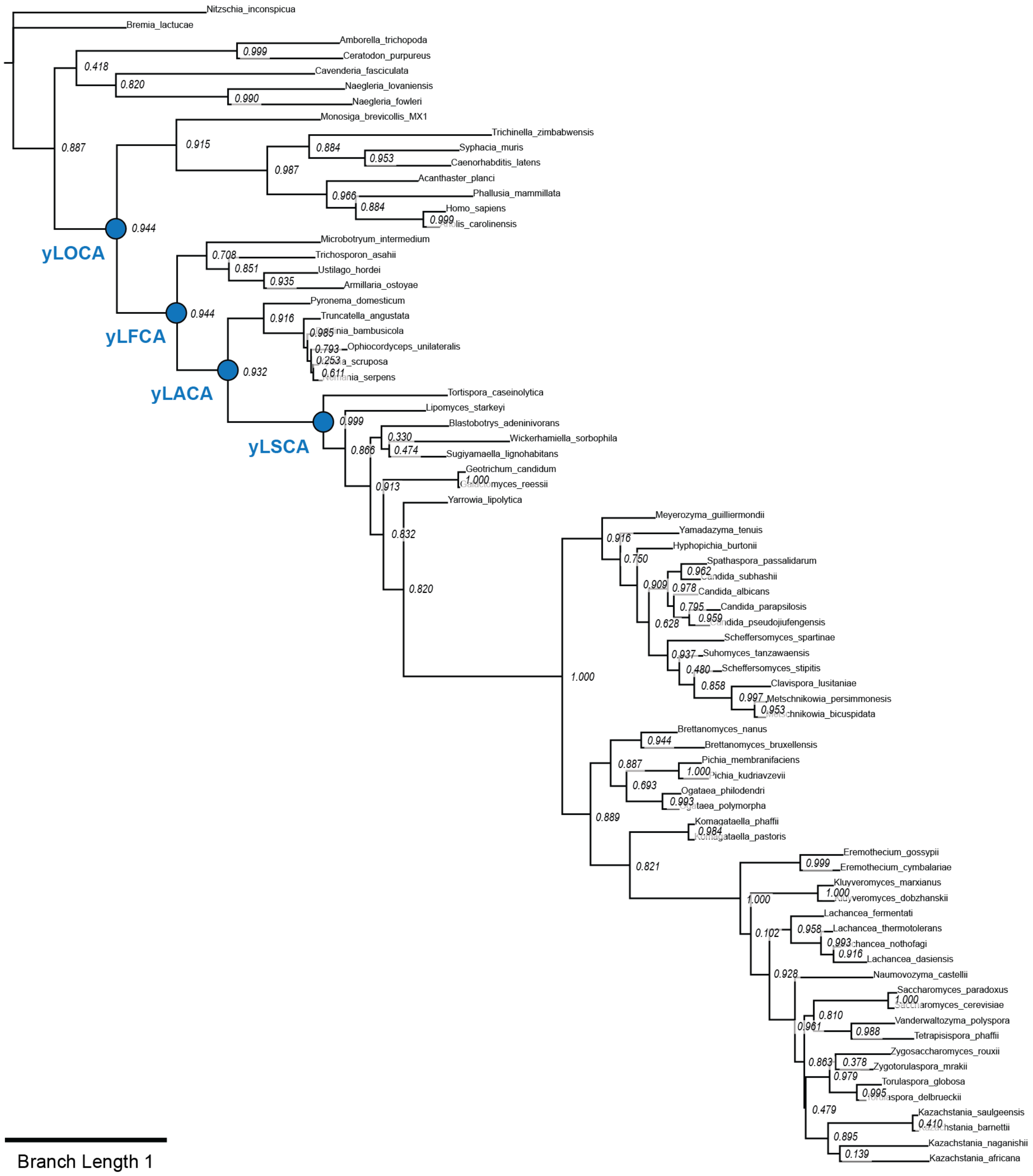


**Figure S4. Yeast-focused phylogenetic tree used for reconstruction of yLSCA, yLACA, yLFCA and yLOCA Mitofilin ancestors.** Reconstructed common ancestors are depicted as blue spheres in the tree. Branch supports are indicated at each node. Branch lengths are shown to scale.

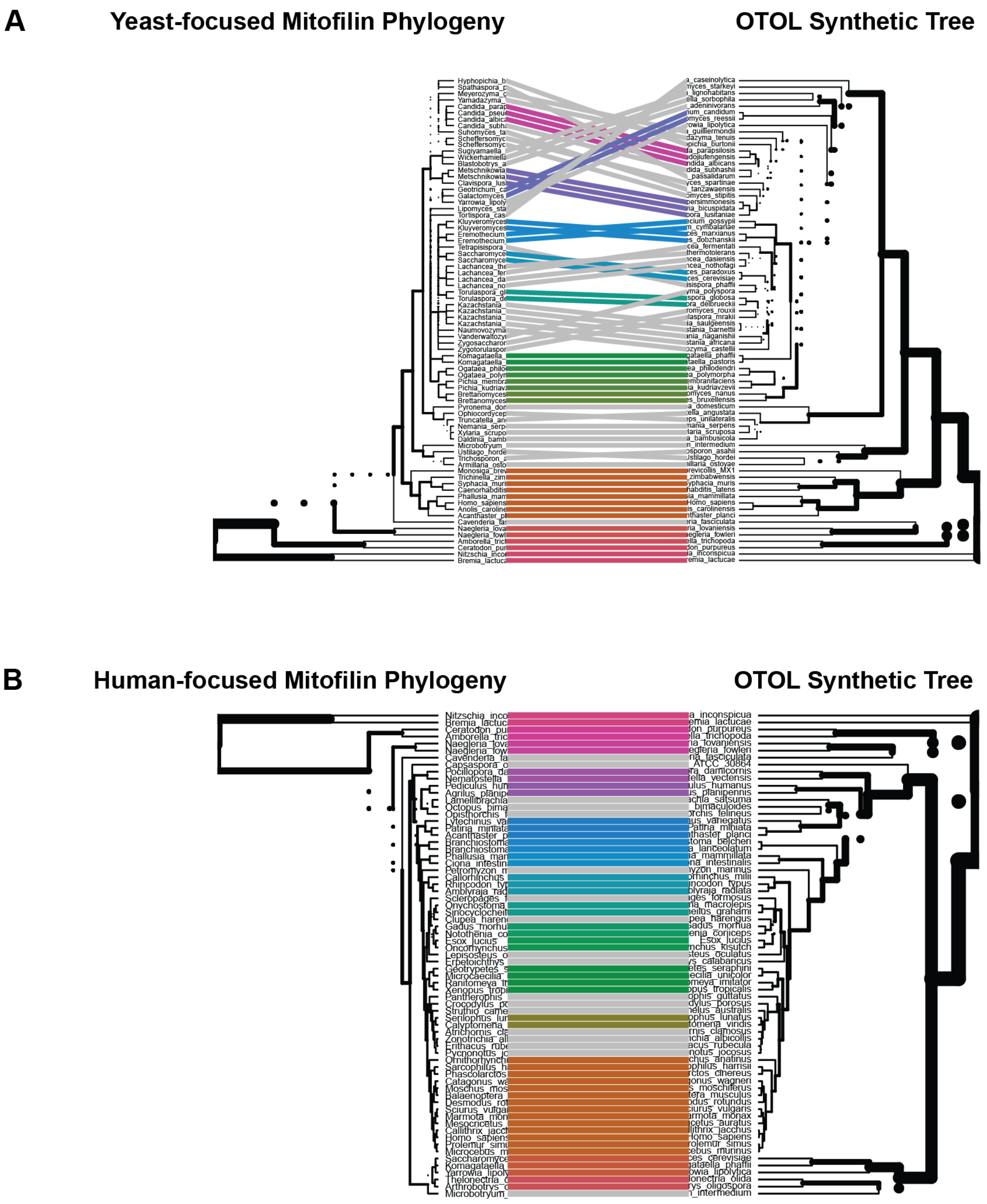


**Figure S5. Tanglegrams of yeast- and animal-focused tree against the Open Tree of Life. (A)** Tanglegram of the yeast-focused tree (left) against the Open Tree of Life Synthetic Tree[5] (right). **(B)** Tanglegram of the animal-focused tree (left) against the Open Tree of Life Synthetic Tree (right).

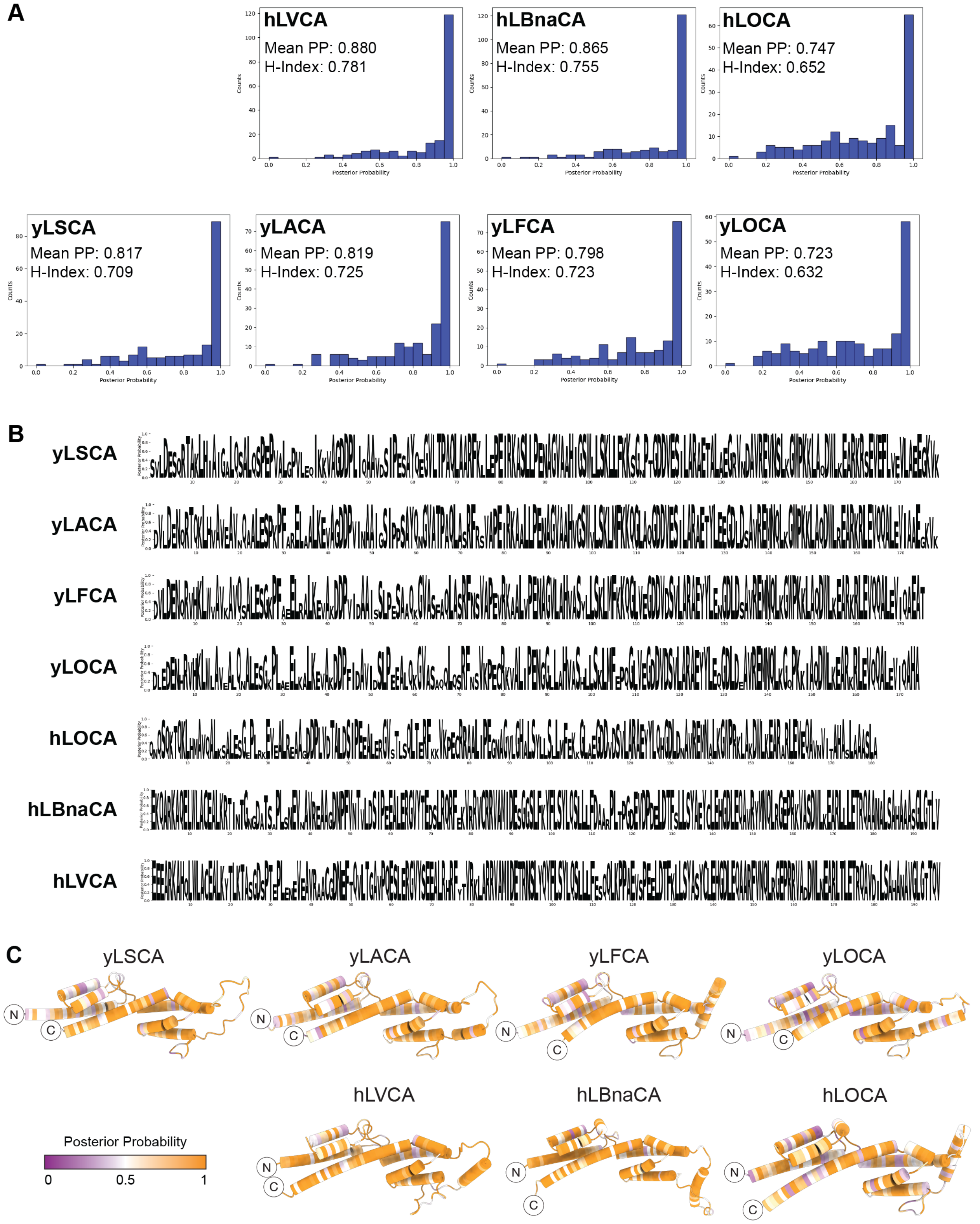


**Figure S6. Support for Ancestral Sequence Reconstruction indicated by posterior probability. (A)** Histograms showing posterior probability (PP) for reconstructed ancestors. The PP for each site is binned in 5% intervals. The H-index indicates the percentage of sites (N%) with a PP larger than N%. **(B)** PP at each reconstructed position across the Mitofilin sequence expressed in WebLogo[6,7] format. The height of each letter corresponds to its PP. **(C)** AlphaFold2[1] models of each reconstructed Mitofilin ancestor with residues colored by PP ranging from low PP (purple) to high PP (orange).

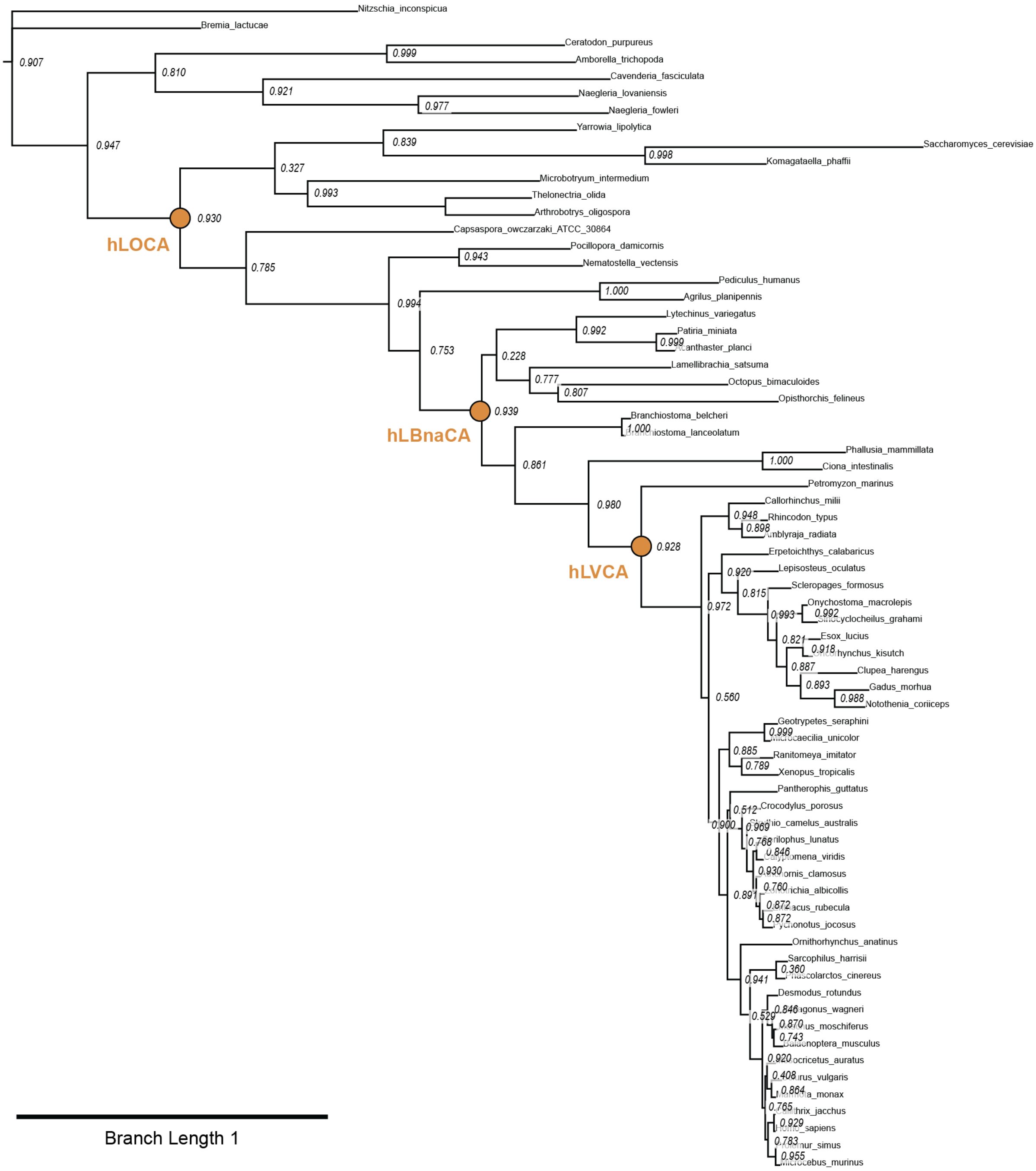


**Figure S7. Animal-focused phylogenetic tree used for reconstruction of hLVCA, hLBnaCA, and hLOCA Mitofilin ancestors.** Reconstructed common ancestors are depicted as golden spheres in the tree. Branch supports are indicated at each node. Branch lengths are shown to scale.

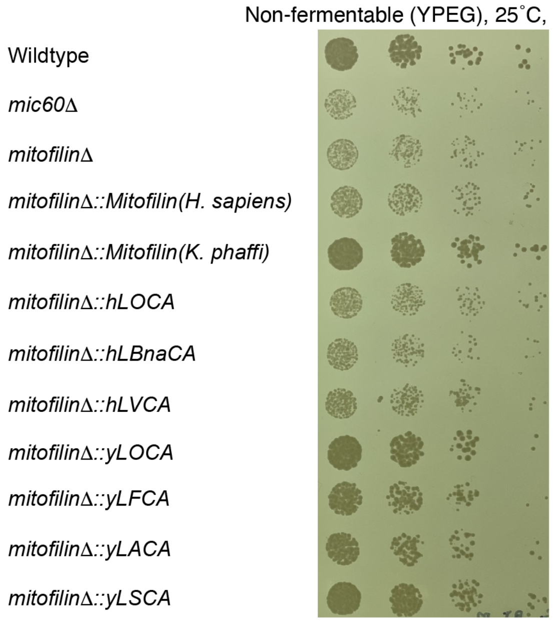


**Figure S8. Mitochondria-dependent growth on non-fermentable media.** Spot assay of yeast Mic60 strains on non-fermentable YPEG agar (1% yeast extract, 2% peptone, 2% glucose) after 5 days at 25˚C. Yeast Mic60 chimeras of all yeast-derived Mitofilin ancestors (yLOCA, yLFCA, yLACA, and yLSCA) and the *K. phaffi* Mitofilin domain grow like wildtype. Yeast Mic60 chimeras of all human-derived Mitofilin ancestors and the *H. sapiens* Mitofilin domain exhibit a knockout-like respiratory growth defect.

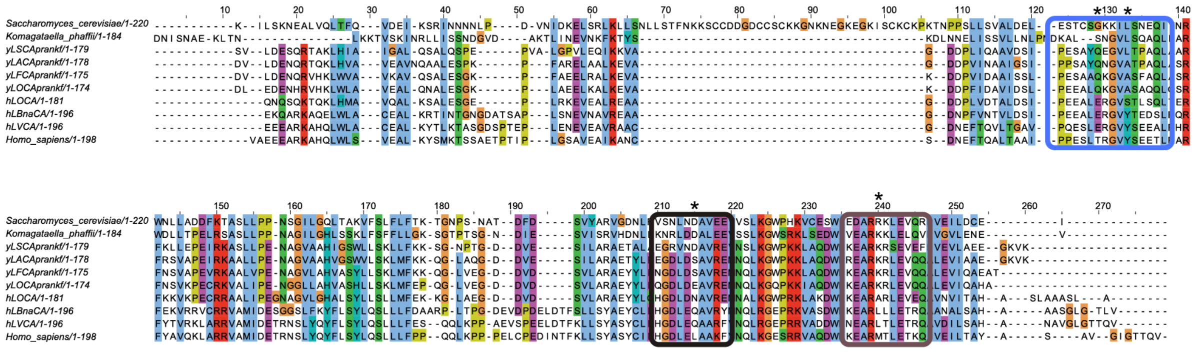

**Figure S9. Multiple sequence alignment highlighting candidate residues explaining functional Mitofilin differences.)** Full-length multiple sequence alignment of Mitofilin domains used for contextualization in Figure 4A. Colored boxes correspond to alignment excerpts displayed in Figure 4A. Candidate residues are indicated by stars (*).

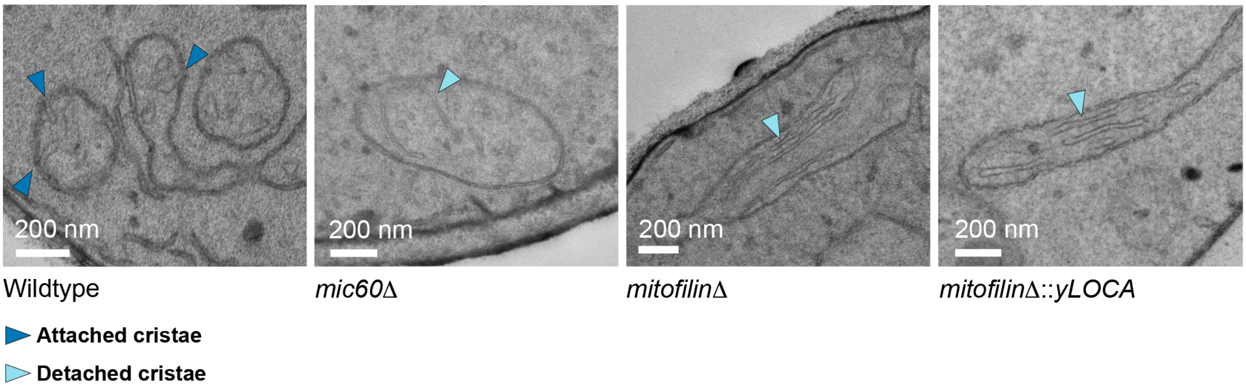


**Figure S10. Transmission electron micrographs of yeast strains grown under respiratory conditions.**
Yeast strains were imaged after 6 hours of respiratory growth in YPEG medium at 30˚C. Arrowheads denote attached (dark blue) and detached (light blue) cristae.

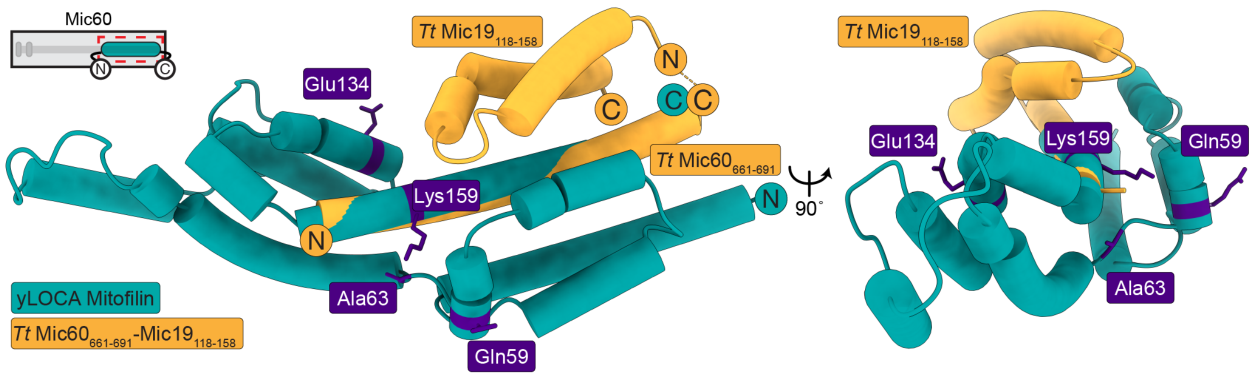


**Figure S11. Mic19-Mic60 interface mapped onto predicted yLOCA Mitofilin structure.** Crystal structure (yellow, PDB 7PV1[8]) of *Thermochaetoides thermophila* Mic60 (residues 661-691; *Tt* $Mic60_{661\text{-}691}$) fused to *Thermochaetoides thermophila* Mic19 (residues 118-158; *Tt* $Mic19_{118\text{-}158}$) superposed onto AlphaFold2 predicted yLOCA structure (teal). The ASR-identified 4-residue signature is depicted as purple residues in yLOCA nomenclature.
